# Variable inbreeding depression may explain associations between the mating system and herbicide resistance

**DOI:** 10.1101/2020.09.25.313940

**Authors:** Megan Van Etten, Anah Soble, Regina S Baucom

**Affiliations:** Biology Department, Pennsylvania State University, Dunmore, PA, 18512; Ecology and Evolutionary Biology Department, 4034 Biological Sciences Building, University of Michigan, Ann Arbor, MI 48109

**Keywords:** Inbreeding depression, herbicide resistance, mating system evolution, crop weeds, *Ipomoea purpurea*, adaptation

## Abstract

Inbreeding depression is a central parameter underlying mating system variation in nature and one that can be altered by environmental stress. Although a variety of systems show that inbreeding depression tends to increase under stressful conditions, we have very little understanding across most organisms how the level of inbreeding depression may change as a result of adaptation to stressors. In this work we examined the potential that inbreeding depression varied among lineages of *Ipomoea purpurea* artificially evolved to exhibit divergent levels of herbicide resistance. We examined inbreeding depression in a variety of fitness-related traits in both the growth chamber and in the field. We paired our examination of inbreeding depression in fitness-related traits with an examination of gene expression changes associated with the level of herbicide resistance, breeding history (inbred or outcrossed), and the interaction of the breeding system and the level of herbicide resistance. We found that, while inbreeding depression was present across many of the traits, lineages artificially selected for increased herbicide resistance often showed no evidence of inbreeding depression in the presence of herbicide, and in fact, showed evidence of outbreeding depression in some traits compared to non-selected control lines and lineages selected for increased herbicide susceptibility. Further, at the transcriptome level, the resistant selection lines had differing patterns of gene expression according to breeding type (inbred vs outcrossed) compared to the control and susceptible selection lines. Our data together indicate that inbreeding depression may be lessened in populations that are adapting to regimes of strong selection.

## Introduction

The plant mating system, often measured as the frequency of self-fertilization, influences evolutionary processes through its impact on effective population size, the distribution of genetic variation among populations, gene flow, rates of homozygosity and heterozygosity among progeny, as well as fitness and the potential to respond to changing and perhaps stressful selective regimes (Barrett & Harder, 2017; Charlesworth & Charlesworth, 1987; Charlesworth & Wright, 2001; Pannell, 2010). The mating system is a highly variable trait among angiosperms, with half of all flowering species reproducing *via* self-pollination at least 20% of the time (Barrett & Eckert, 1990). Within mixed mating species, populations display a striking amount of variation in outcrossing rates (Whitehead et al., 2018). However, the underlying causes of such variation across populations and the extent to which environmental factors underlie this variation is poorly understood. In fact, parameters that are central to understanding plant mating system diversity in nature (Goodwillie et al., 2005) such as pollen limitation (Harder & Aizen, 2010), inbreeding depression (Cheptou & Donohue, 2011), reproductive assurance (Kalisz et al., 2004) and pollen discounting (Kohn & Barrett, 1994) each strongly depend on environmental conditions, but how such conditions and environmental changes may alter these mating system parameters is broadly unknown.

Thus far, current data across a range of plants suggest anthropogenic modifications of the environment--e.g., forest fragmentation, urbanization--are correlated with higher rates of selfing and biparental inbreeding (Cheptou & Avendaño, 2006; Eckert et al., 2010), but the underlying causes of these correlations are unclear. In many cases, it is likely that the increased selfing rate represents plastic rather than evolved changes to the mating system. However, there are a handful of studies that suggest that the mating system can evolve due to human-mediated changes in the environment. For example, experimental populations of *Mimulus guttatus* reared in the absence of pollinators over five generations led to the evolution of floral characters associated with self-pollination, showing that the disruption of pollinator services can select for selfing types (Bodbyl Roels & Kelly, 2011). Similarly, plants adapted to heavy metals often, but not always, exhibit higher rates of self-pollination compared to non-tolerant plants (Antonovics et al., 1971; Antonovics, 1968; Cuguen et al., 1989; Dulya & Mikryukov, 2016; Lefebvre, 1970; Mousset et al., 2016). In another example, variation in the selfing rate across populations of *Ipomoea purpurea*, the common morning glory, was strongly correlated to the level of herbicide resistance, with the most resistant populations exhibiting higher rates of selfing (Kuester et al., 2017).

For the maintenance of such an association between higher selfing and adaptation, the benefit of being both environmentally adapted and highly selfing would have to outweigh the costs associated with selfing -- namely, fitness costs incurred from inbreeding depression (Levin, 2012). The cost of selfing may be further amplified by making it more difficult for selfed progeny to adapt to novel environments. In fact, theoretical work suggests that increased selfing in populations adapting to heterogeneous environments may be impeded by inbreeding depression, thus leading to higher rates of random mating (Epinat & Lenormand, 2009).

Unfortunately, we have little to no empirical understanding of how adaptation to an environmental stressor may be influenced by inbreeding depression in plants (or *vice versa*). Work examining inbreeding depression writ large in plant systems as well as the interaction of inbreeding depression and herbivory shows that typically, inbreeding depression is more pronounced in ‘stressful’ or novel environments (Fox & Reed, 2011) and that inbreeding reduces resistance and tolerance to herbivores (Carr & Eubanks, 2002; Du et al., 2008; Ferrari et al., 2007; Hull-Sanders & Eubanks, 2005; Ivey & Carr, 2012). However, beyond the plant-herbivore work, there is little known about inbreeding depression in plants resistant or tolerant to stressors, meaning that a significant gap remains in our understanding of how inbreeding depression may influence mating system evolution and especially so in scenarios of plant adaptation to strong, human-mediated regimes of selection.

Here, we examined the potential for inbreeding depression in genetic lineages of *Ipomoea purpurea* selected for divergent levels of herbicide resistance to make inferences about the previously identified association between the mating system and resistance. Using individuals produced by inbreeding and outcrossing, we examined phenotypic traits and transcriptomic responses to answer: (1) Do lines that vary in their level of resistance show divergent levels of inbreeding depression in post-herbicide survival and biomass traits? (2) Is there evidence of inbreeding depression in fitness-related traits (flowering time, flower number, seed number, and seed germination) in lines that vary for resistance in field conditions, in both the absence and the presence of herbicide? 3) At the transcriptome level, are there differences in gene expression associated with breeding history, resistance, or the interaction between breeding history and resistance?

## Methods

### Study species

The common morning glory, *Ipomoea purpurea*, is a common weed of agricultural fields as well as other sites of disturbance. It provides an excellent study system with which to examine the influence of inbreeding depression on the evolution of the mating system. This mixed-mating species displays an average outcrossing rate of 0.5 (range across 24 natural populations: 0.2-0.8) (Kuester et al., 2017), and exhibits inbreeding depression in a variety of mating-system (Chang & Rausher, 1999) and growth-related traits (Darwin, 1876). Additionally, *I. purpurea* populations exhibit variation in resistance to glyphosate, which is the active ingredient in the herbicide RoundUP (Kuester et al., 2015), and this variation in glyphosate resistance among populations is negatively correlated with the mating system, such that the most resistant populations display the lowest outcrossing rate (Kuester et al., 2017).

### Generation of experimental seeds

We previously generated artificially evolved selection lines that varied in the level of glyphosate resistance and used these lines to perform the current study (Debban et al., 2015). Selection lines were generated using *I. purpurea* maternal individuals from the same initial base population (full design described in (Debban et al., 2015)), and after three generations of artificial selection using two replicated selection lines per direction of selection (*i*.*e*., 2 independent replicates of crosses used for three directions of selection -- increased resistance (R), decreased resistance (S), and control (C) selection lines), the increased resistance selection lines exhibited ∼75% survival post-herbicide application in the field whereas ∼55% and ∼45% of individuals from the (non-selected) control and susceptible (decreased resistance) lines survived, respectively (Debban et al., 2015). For the work described here, we chose the 3 most/least resistant families from the Debban et al (2015) field screening from each of the two replicated R and S selection lines, respectively, for a total of 6 families per direction of selection. We generated seeds by crossing maternal individuals from the second generation of artificial selection from each chosen family to three paternal individuals from each respective direction of selection regardless of replication line (hereafter outcrossed seeds) and likewise generated self-pollinated seeds (hereafter inbred seeds) from each maternal individual (Fig 1A). Parents for the control lines were randomly chosen but following the same general design. This led to the production of outcrossed and inbred seeds for six families per direction of selection (hereafter selection lines), and thus a total of 18 families for use in the growth chamber and field experiments.

**Fig 1.**
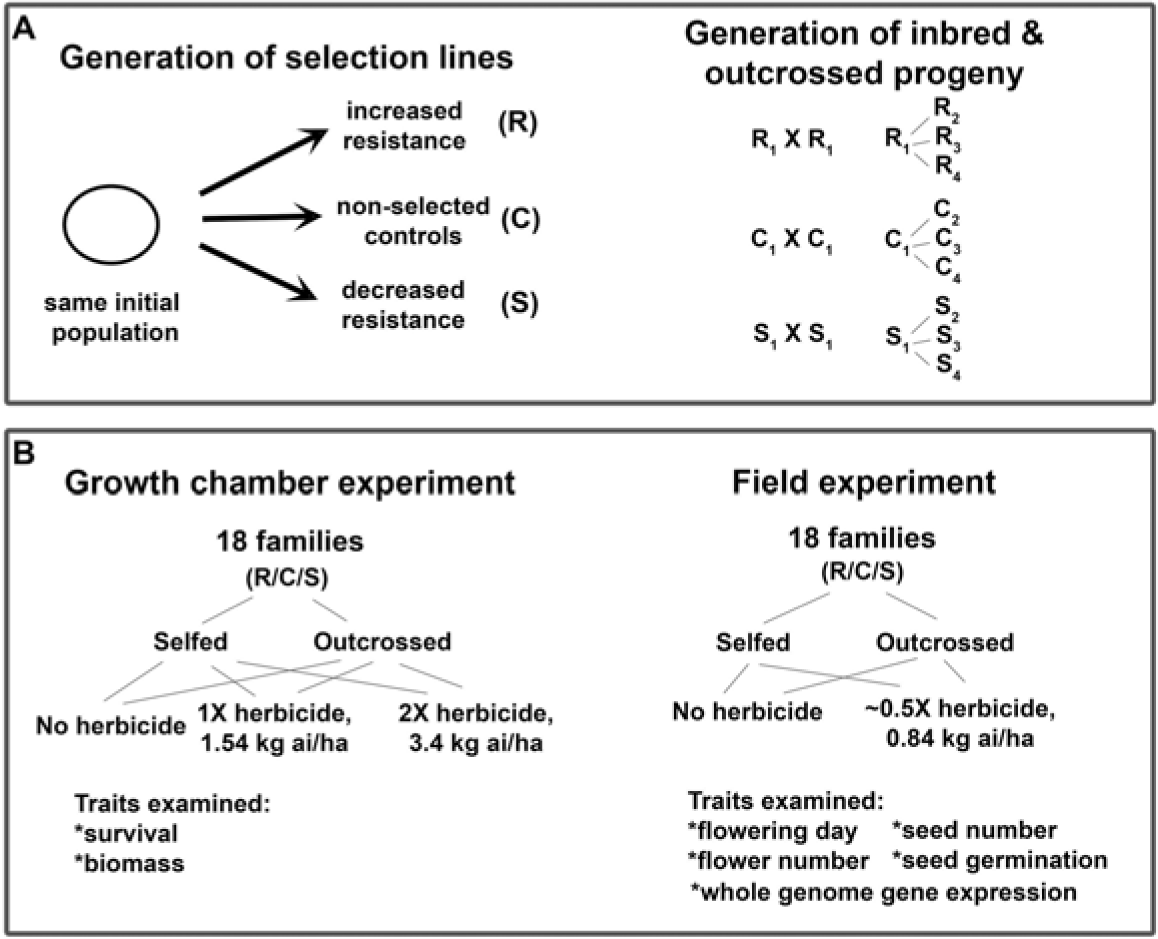
Experimental design used in this work. A) We performed three generations of artificial selection for both increased (R) and decreased (S) resistance, as described in (Debban et al., 2015), and used the same crossing design to generate non-selected controls (C). After two generations of selection, we selfed and outcrossed 18 maternal families (6 each direction of selection) to generate experimental seeds. B) We performed a growth chamber experiment to examine inbreeding depression in survival and biomass remaining post-herbicide. We paired our analysis of inbreeding depression in fitness traits in the field with a transcriptome survey to examine potential changes in gene expression associated with direction of selection (R/C/S), breeding status (I/O) and the interaction of direction of selection and breeding status (i.e., I/O within R/C/S).

#### Growth chamber & field experiments

We performed a growth chamber and a field experiment to examine the potential that inbreeding depression varied according to resistance status and in response to herbicide exposure. In the growth chamber experiment, we focused on plant biomass and measures of damage post-herbicide application whereas in the field we examined estimates of fitness--flowering day, flower number, seed number, and germination of seeds produced (see Fig 1 for experimental design).

### Growth chamber experiment

On April 4, 2019, we planted five replicates of both outcrossed and inbred seeds of each family in Cone-Tainers (Stewe and Sons) in each of three treatment environments--two doses of glyphosate (1.54 kg ai/ha, 3.4 kg ai/ha) and a no-herbicide control. The lower dose of herbicide is the suggested field dose (hereafter 1X) whereas the higher dose is slightly less than two times the field dose (hereafter 2X). We elected to use two levels of herbicide in this experiment since assays of natural populations have shown high survival in a few populations at 3.4 kg ai/ha. In total we assayed 855 plants. Plants were allowed to grow for 48 days until the 8-leaf stage at which point herbicide was applied using a CO_2_-pressurized handheld sprayer (R and D Sprayers, Opelousas, LA). Prior to herbicide application, plant height (cm) and number of leaves were recorded. Fifteen days post-herbicide, we recorded death as a metric trait (dead/almost dead or green and healthy), and recorded height (cm). We then harvested each plant in the experiment, dried the plants for 3 days at 70C, and weighed each for an estimate of dried above-ground biomass.

### Field experiment

We planted five replicate seeds from each maternal and paternal line combination into two blocks and two herbicide treatments (no herbicide control, herbicide) in a randomized block design at Matthaei Botanical Gardens (University of Michigan) in the spring of 2015. Thus, within each block there were 5 selfed replicates and 15 outcrossed replicates (5 from each father) resulting in 1440 plants total. Seeds were scarified and planted with a 1 m separation. After approximately 6 weeks of growth, we applied 0.84 kg ai/ha glyphosate using a CO_2_ sprayer to plants in the herbicide treatment. We elected to apply a low level of herbicide at this later stage of growth since we wanted to stress the plants with the herbicide but not cause a significant amount of death. Throughout the experiment, plants were watered as needed to prevent wilting, and we mowed the field once to reduce competition.

We measured plant height (cm) at six weeks, just prior to spraying. Two weeks after herbicide application, we measured the height of the plants. We recorded flower number twice weekly on each plant about 1 week after plants started flowering. Once seeds developed, we sampled them from three rounds of collection (between September 20 - October 10): in the first round we collected all developed seeds from plants in the control treatment, as seeds on these plants matured faster than those from the herbicide treatment, and in the second and third round we collected all plants in brown paper bags and then separated seeds from the plants, noting which seeds were developed or still green.

Finally, we examined the potential that seed germination differed according to herbicide treatment, selection direction, and breeding type since previous work has identified a cost associated with herbicide resistance in this species in terms of low germination of resistant individuals (Van Etten et al., 2016). To do so we performed a germination assay using petri dishes in the lab. We placed five randomly chosen seeds from each field plant in a petri dish with 10 ml water and assessed germination at three, five, seven, and 14 days post water exposure. At day 14 we scarified any seeds that had not germinated and assessed germination once more to determine if some seeds were still viable but needed scarification to germinate. Total germination was considered the number that germinated both prior to and post scarification.

### Data analysis

All analyses were performed with the R programming language (Team, 2015) in RStudio (vers. 1.3.959). We used logistic regression and mixed model ANOVAs to examine the response to artificial selection in the growth chamber experiment and mixed model ANOVAs to determine if there were significant differences in the expression of inbreeding depression in both the growth chamber and field experiments. Using data from the growth chamber experiment, we assessed the potential for response to artificial selection by testing for a main effect of selection direction (R/S/C) on death and dry biomass. We used a generalized linear model (glm function; Team R, 2015) with death score as the dependent variable and selection line (R/S/C), breeding type (I/O), maternal line nested within selection line, and treatment as the independent variables and specified a logistic link function. Because this model would not converge, we elected to use a simpler model by removing the maternal line and interaction effects from the final model (death score ∼ selection line (R/S/C) + breeding type (I/O) + treatment). We assessed significance of variables by comparing analysis of deviance after removing variables from subsequent models using the anova() function.

To examine biomass, we performed a mixed model ANOVA using the lmer function (Bates et al., 2015), with biomass as the dependent variable and selection line (R/S/C), breeding type (I/O), and treatment (1X vs 2X herbicide) as fixed independent variables. To control for differences in plant size, biomass was standardized by dividing each individual biomass value, in both herbicide treatments, by the average biomass of each respective maternal line from the control environment. Maternal line nested within selection line was included as a random, independent variable in the final model and biomass was log transformed prior to analysis to improve the fit of the residuals. We ran a preliminary analysis including all interactions between breeding type, treatment, and selection line, but elected to reduce the final model using the step() function in the lmerTest package (Kuznetsova et al., 2017) since our primary interest was in the main effects and the potential for the interaction between selection line and breeding type (I/O). Our final model was biomass ∼ selection line (R/S/C) + breeding type (I/O) + treatment + maternal line (nested within selection line) + breeding type*treatment + breeding type*selection line. Further, we elected to perform analyses within treatment (1X vs 2X dose) separately because we uncovered a significant breeding type by treatment interaction and because the overall model showed a very significant treatment effect (*i*.*e*., a decline in biomass given a higher dose of herbicide) which was neither surprising nor a main interest in this study.

We next examined fitness related traits (day of first flower (flowering day), flower number, seed number, and seed germination) in the field to determine if there were differences in the level of inbreeding depression associated with selection line (R/S/C) and in response to stress from the herbicide. We first performed mixed model analyses using the lmer function of lme4 (Bates et al., 2015) to determine if herbicide application altered day of flowering, flower number, and total seed number. We log transformed dependent variables and used the following general model for both traits separately: trait ∼ treatment + block + breeding type (I/O) + selection line (R/S/C) + maternal line (nested within selection line) + breeding type*selection line + treatment*breeding type + treatment*selection line + breeding type*selection line*treatment. We then restricted further analyses to within treatment using a reduced general model (trait ∼ block + breeding type (I/O) + selection line (R/S/C) + maternal line (nested within selection line) + breeding type*selection line). In each model, maternal line was a random factor and was nested within selection line (R/S/C). To examine the effect of breeding history and level of resistance on seed germination, we used a generalized linear model to perform logistic regression with the proportion seeds that germinated as the dependent variable and breeding type (I/O), selection line, the interaction of breeding type and selection line, and block as independent variables. We did not assess the potential for maternal line variation in seed germination as preliminary logistic regressions using maternal line as a random factor would not converge. Thus, we cannot evaluate the potential for genetic variation underlying germination in this work. As above, we first included a treatment factor in the logistic regressions but focus reporting our results within the two environments (herbicide, no herbicide) separately.

### Inbreeding depression

In both the field and growth chamber experiments, we calculate inbreeding depression (δ) for traits for each maternal line within direction of selection (R/S/C) and treatment using the formula δ□= □ (1−trait selfed plants/trait outcross plants).

#### RNAseq experiment

### Tissue collection

We collected leaf material from a subset of the families grown in the field experiment to examine the potential for gene expression changes associated with artificial selection and the potential for inbreeding depression. Because we were explicitly interested in understanding gene expression differences associated with inbreeding depression, we sampled leaf material from two families per selection line that exhibited either high or low inbreeding depression, based on the first height measurement (i.e., families that showed little or no IBD in height vs families that showed significant difference in the height of inbred individuals versus outcrossed individuals). We sampled leaf tissue from a total of four families per selection line (R/S/C) and at two time points post-herbicide application--eight and 32 hours after spraying with herbicide. Within each time point, we randomly chose one individual per family*selfing treatment*spray treatment and froze 1-2 young leaves of individuals in liquid nitrogen (a total of 96 individuals).

### RNA extraction and library preparation

We extracted RNA using the Qiagen RNeasy plant kit and constructed Individually indexed libraries using the Illumina TRUseq96 indexer mRNA stranded kit. We pooled the resulting libraries and ran on 7 lanes of 50 Single end sequencing on the Illumina HiSeq 4000. This resulted in 88 GB of data and on average 28 million reads per individual.

### Sequencing and data processing

We removed adaptors using cutadapt (Martin, 2011), with a maximum error rate of 10% and discarded sequences < 21 bp. The reads from all individuals were digitally normalized using khmer (Crusoe et al., 2015), with a kmer cutoff of 20 and a kmer size of 23. We assembled transcripts using Velvet (Zerbino, 2010) through the Oases pipeline (Schulz et al., 2012), which creates assemblies over a range of k-mer lengths (from 21-61 in increments of 10) and then merges the results into a final assembly. We removed resulting contigs less than 100 bp long. We then used LACE (Davidson et al., 2017) and CORSET (Davidson & Oshlack, 2014) to obtain the SuperTranscript from the Oases transcriptome as well as obtain gene level counts for each individual.

### Annotating the transcriptome

We annotated SuperTranscripts using a tblastx (default settings; (Altschul et al., 1990)) against the *Arabidopsis thaliana* cDNA (release 10) and an annotated *Ipomoea nil* transcriptome (Hoshino et al., 2016). We retained the best match (lowest e-value) for each gene assuming it had an e-value of at least 0.001 and a length of more than 10 nt.

### Data analysis

We used network analysis to identify groups of genes whose expression differed according to selection line, breeding type, and breeding type within selection line. We performed comparisons using WGCNA (Langfelder & Horvath, 2008), or weighted correlation network analysis, which allowed us to find clusters of correlated genes associated with each of our main comparisons of interest while reducing the dimensionality of the data. In brief, WGCNA creates groups of genes, called modules, based on pairwise correlated expression. The expression of the resultant eigengene can then be correlated with a factor (*e*.*g*., selection line, or breeding type, etc.) to determine if the module’s expression levels correlate with a comparison of interest.

The set of transcriptomic data used in WGCNA depended on the comparison of interest. To examine which genes had differential expression between selection lines, we first broke the data into three treatment groups: 1) the non-sprayed treatment, and plants that were sprayed with herbicide and sampled at 2) eight and 3) 32-hours post-spray. A preliminary principle components analysis showed large differences in expression between these three treatment groups, suggesting that differential expression patterns may differ between the three treatment groups and should be considered separately. Thus, for each of these groups, we created modules then determined which eigengenes correlated with the selection line (where selection lines were given a value of S = 1, C = 2, R = 3). This analysis would identify eigengenes with opposite expression in the R and S lines and an intermediate expression in the C lines. To examine overall differences due to breeding type, we again created modules separately for the non-sprayed, sprayed 8-hour and 32-hour groups, then determined which eigengenes were correlated with breeding type (outbred = 1, inbred = 2). This analysis would identify eigengenes with differing expression between inbred and outcrossed individuals within each of the environments. Finally, to examine whether there were differences between lines in the genes that were affected by breeding type, we used a consensus analysis for each of the 3 groups (non-sprayed, sprayed 8-hour and sprayed 32-hour). In this analysis, modules are created using the whole group, but the correlation between eigengenes and treatment is done separately for each subgroup (in this case the selection lines). This analysis would identify eigengenes with expression differences between inbred and outcrossed individuals within each of the selection lines.

For all of the above analyses, we chose to use a signed analysis since we were interested in capturing the direction of expression differences. The optimal soft thresholding value differed depending on the analysis and was determined as suggested by the program creators. The genes in modules that were significantly correlated with the treatment of interest were identified and their GO term enrichment determined using ShinyGO (Ge et al., 2020) based on the *Arabidopsis thaliana* annotation (P-value cutoff = 0.05 except for saddlebrown module for spray, 8 hour: P-value 0.2; 30 most significant terms were kept).

## Results

### Growth chamber experiment shows differences in death and biomass between artificially evolved selection lines and according to breeding history

We examined response to two different herbicide doses -- both a recommended field dose (1X) and approximately two times the recommended dose (2X) -- in the growth chamber to capture potential differences between selection lines and breeding types (inbred and outbred plants) at early growth stages and in controlled conditions.

Plant death differed between selection lines, breeding types, and herbicide treatment. We found a significant effect of selection line (Chi-square = 29.89, p-value < 0.001; Table S1) such that individuals in the resistant selection line exhibited significantly lower proportion death than individuals from the control (45% R vs 60% C: Chi-square = 5.90, p-value 0.015) and susceptible (45% R vs 70% S: Chi-square = 24.57, p-value < 0.001) selection lines. The difference in proportion death between susceptible and control lines was marginally significant (Chi-square = 3.21, p-value = 0.073). We found a highly significant treatment effect (1X vs 2X herbicide dose; Chi-square = 62.54, p-value < 0.001; Table S1) with 43% and 75% individuals being either dead or almost dead in the 1X and 2X herbicide dose, respectively. We additionally found a significant effect of breeding type (Breeding type effect, Chi-square = 4.96, p-value = 0.026; Table S1), with individuals produced by inbreeding exhibiting higher proportion death (66%) compared to individuals produced by outcrossing (57%) in the presence of herbicide (Table S1). This effect appeared to be driven predominantly by the control lines (Breeding type effect, Chi-square = 5.67, p-value = 0.017) rather than the resistant (Breeding type effect, Chi-square = 0.467, p-value = 0.494) or susceptible lines (Breeding type effect, Chi-square = 0.888, p-value = 0.346).

Dried biomass post-herbicide also differed according to selection line, breeding type, and herbicide treatment. As expected, we found a significant effect of herbicide dose (Treatment effect, F-value = 14.82, p-value < 0.001; Table S2) such that the biomass of individuals was reduced by 18% in the 2X compared to the 1X dose. Because of this large difference, we analyzed the two treatments separately.

We uncovered somewhat different dynamics for dried biomass when exposed to different herbicide rates. We found a significant effect of breeding type at the 1X herbicide dose, with inbred individuals exhibiting significantly lower biomass compared to outbred individuals (Least square means biomass (±se): 0.35 g (±0.03) inbred vs 0.42 g (±0.02) outcrossed individuals). This general trend was apparent across both the control and susceptible individuals, but not individuals from the resistant selection lines (Fig 2B). Interestingly, at the 2X dose, we found a significant interaction between selection line and breeding type (F-value = 4.59, p-value = 0.011; Table S2): while inbred individuals from the control and susceptible selection lines exhibited lower biomass compared to outcrossed individuals (∼27% (C) and ∼15% (S) lower biomass of inbred compared to outbred individuals, Fig 2B), in the resistant selection line, the inbred individuals exhibited significantly *greater* biomass than individuals produced by outcrossing (LSMeans (±se): 0.41 g (±0.04) inbred vs 0.34 g (±0.03) outcrossed). Thus, at this high dose of herbicide, inbred resistant individuals produced 23% greater biomass compared to outcrossed resistant individuals. This suggests that in terms of post-herbicide biomass, resistant individuals exhibited no evidence of inbreeding depression and instead showed the opposite pattern (*i*.*e*. outbreeding depression).

**Fig. 2.**
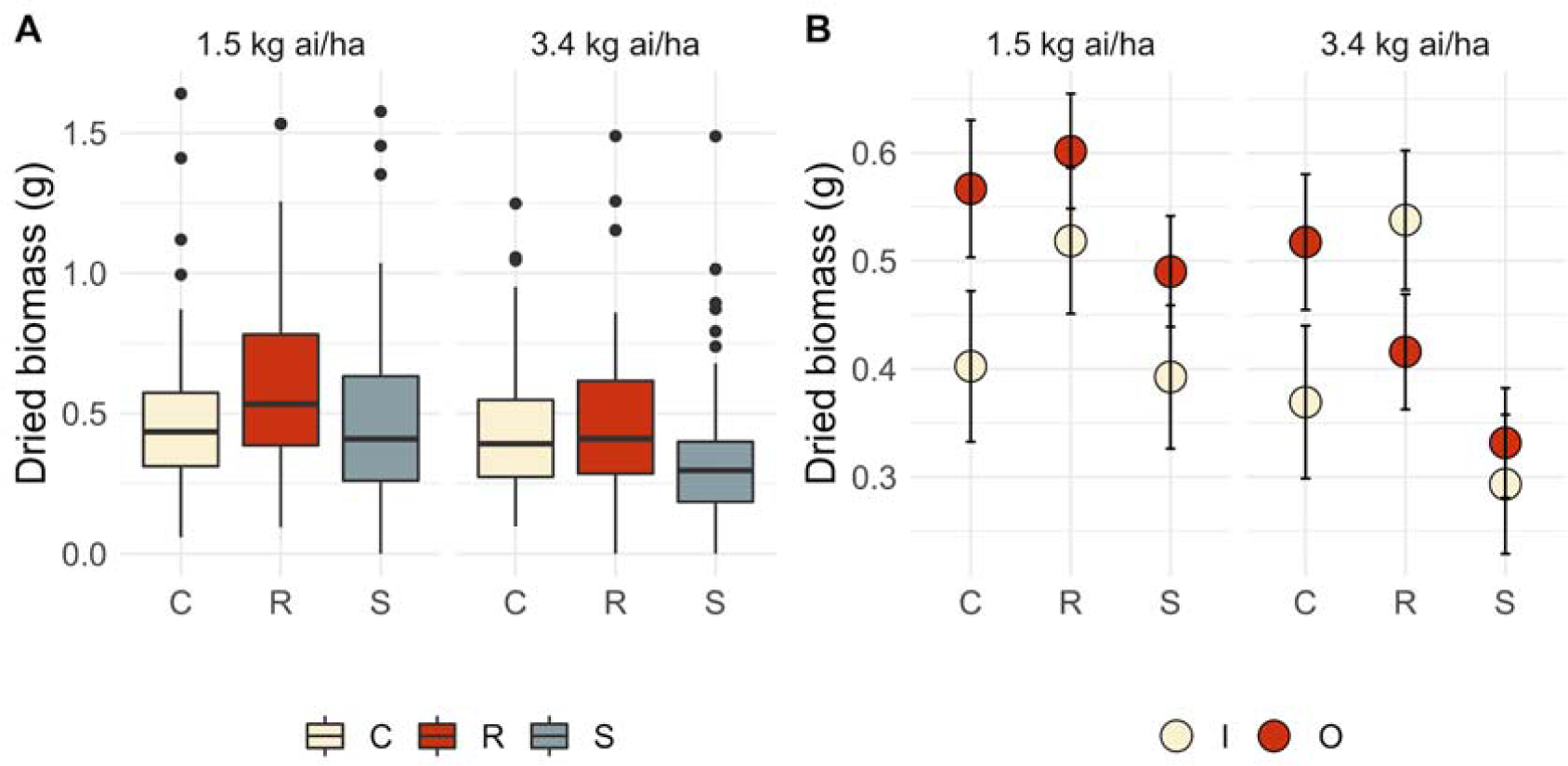
Dried biomass of individuals per A) selection line at the field dose (1.5 kg ai/ha) and approximately two times the field dose (3.0 kg ai/ha) and B) averaged according to selection line and breeding status (I/O) from the growth chamber experiment. Dried biomass for each individual was standardized by average biomass per family from the non-herbicide control environment.

### Field experiment shows differences among selection lines for inbreeding depression

We examined fitness-related traits in field conditions to determine if the selection lines showed differences in the level of inbreeding depression in either the absence or presence of herbicide.

We applied a low rate of herbicide to the plants after six weeks of growth (see Methods), which was not a strong enough rate to cause death (0 plants died in the herbicide present environment). However, this application rate significantly influenced fitness. We found a highly significant effect of herbicide application on flowering day (Treatment effect, F-value = 8058.05, p-value < 0.001), the number of seeds produced (Treatment effect, F-value = 1248.48, p-value < 0.001), and seed germination of seeds produced by experimental individuals (Treatment effect, Chi-square = 390.39, p-value < 0.001). The day of first flowering was delayed by ∼30 days in the herbicide compared to the control environment, which led to an ∼89% reduction in the overall number of seeds produced between control and sprayed plants. Germination was likewise reduced, from an average of 62.7% among seeds produced in the control environment to 30.9% among seeds produced in the sprayed environment (Table S3). We found no evidence for a significant difference in the timing of first flowering, the total number of flowers, the number of seeds produced, or in seed germination between the selection lines, either in the presence or absence of herbicide (Table S4) indicating that fitness did not differ between selection lines at this low dose of herbicide.

We did, however, uncover evidence for inbreeding depression in fitness estimates in both the control and herbicide present environments (Table S4). In the control environment, we found that individuals produced by inbreeding flowered later (0.77 days later, on average; Breeding effect, F-value = 8.64, p-value = 0.003; Table S4) and produced fewer seeds (LSMeans (±se): 2245 (±115.5) inbred vs 2598 (±69.5) outcrossed seeds; Breeding effect, F-value = 5.28, p-value = 0.02; Table S4), and that seeds exhibited lower germination compared to individuals produced by outcrossing (59% inbred vs 64% outcrossed seeds; Breeding effect, F-value = 5.45, p-value = 0.02; Table S4). Both the control and susceptible lines, but not the resistant lines, exhibited inbreeding depression for the number of seeds produced in the absence of herbicide (Fig 3A).

**Fig. 3.**
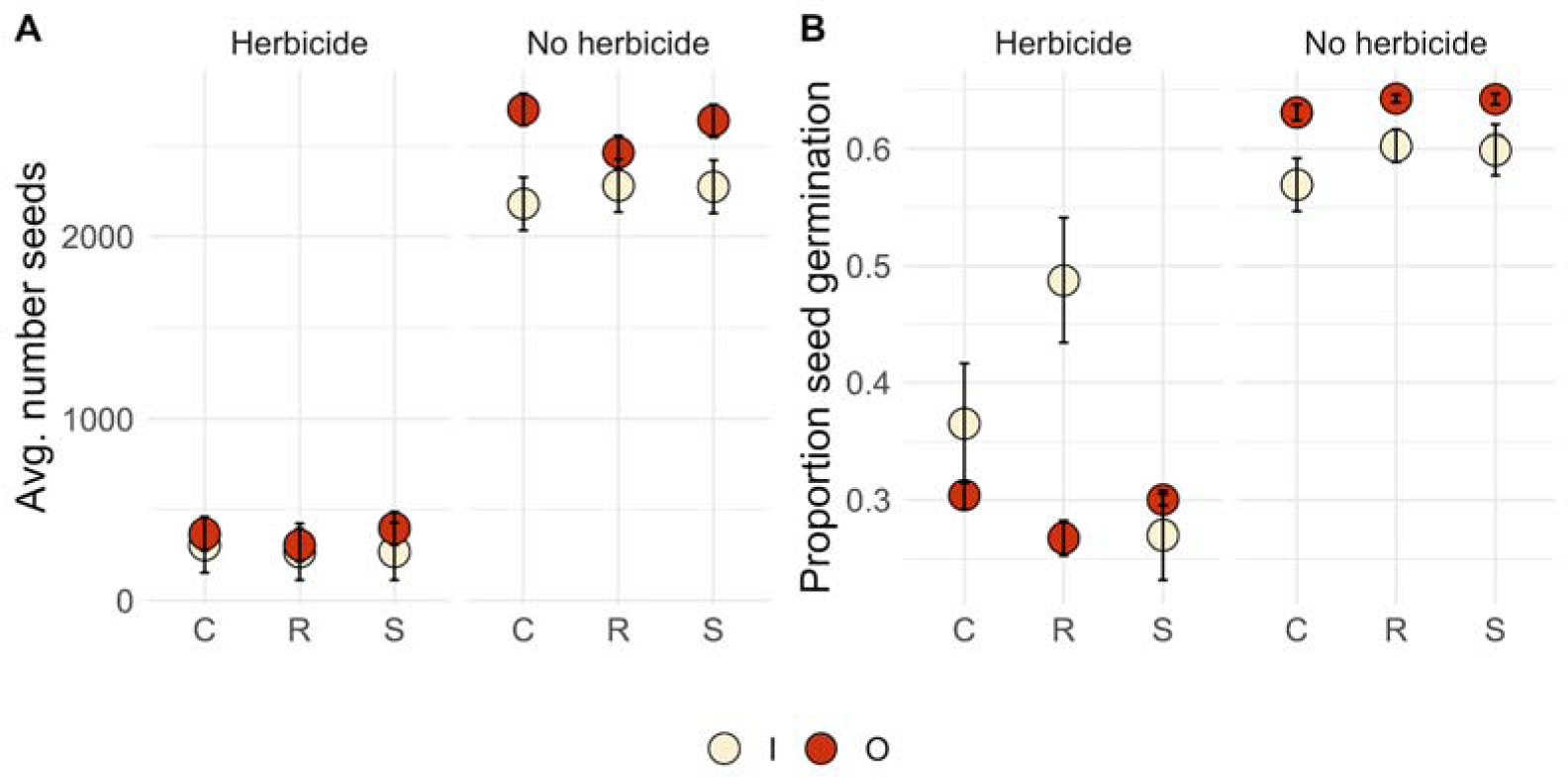
The average number of A) seeds and B) proportion seed germination in both the presence and absence of herbicide in the field according to breeding status (I/O) and selection line. Dots show least square means (± standard error).

In the presence of herbicide, individuals produced by inbreeding produced fewer flowers (LSMeans (±se): 8.46 (±0.47) inbred vs 10.24 (±0.30) outcrossed; Breeding effect, F-value = 12.03, p-value < 0.001; Table S4) and seed (LSMeans (±se): 274 (±42.9) inbred vs 303 (±28.1) outcrossed; Breeding effect, F-value = 6.14, p-value = 0.01; Table S4). Strikingly, we found a significant interaction between breeding type and resistance line (F-value = 6.73, p-value = 0.04; Table S4; Fig 3B) for proportion seed germination. In the herbicide present environment, 49% of the seeds produced by resistant, inbred individuals germinated whereas 27% of the seeds produced by resistant, outcrossed individuals germinated (Table S5). This differed from both the control and susceptible lines, which showed similar seed germination between inbred and outcrossed individuals (Table S5).

Thus, although there was no indication that fitness varied between selection lines at this low application rate, the level of inbreeding depression appeared to vary between selection lines in the presence of herbicide. Across both experiments, the general pattern is that control and susceptible lines exhibited evidence for inbreeding depression in biomass and seed production in the presence or absence of herbicide (Fig 4 A-C), whereas the resistant lines showed no evidence of inbreeding depression (seed number, Fig 4B) or evidence of outbreeding depression (biomass, seed germination; Fig. 4 A,C).

**Fig 4.**
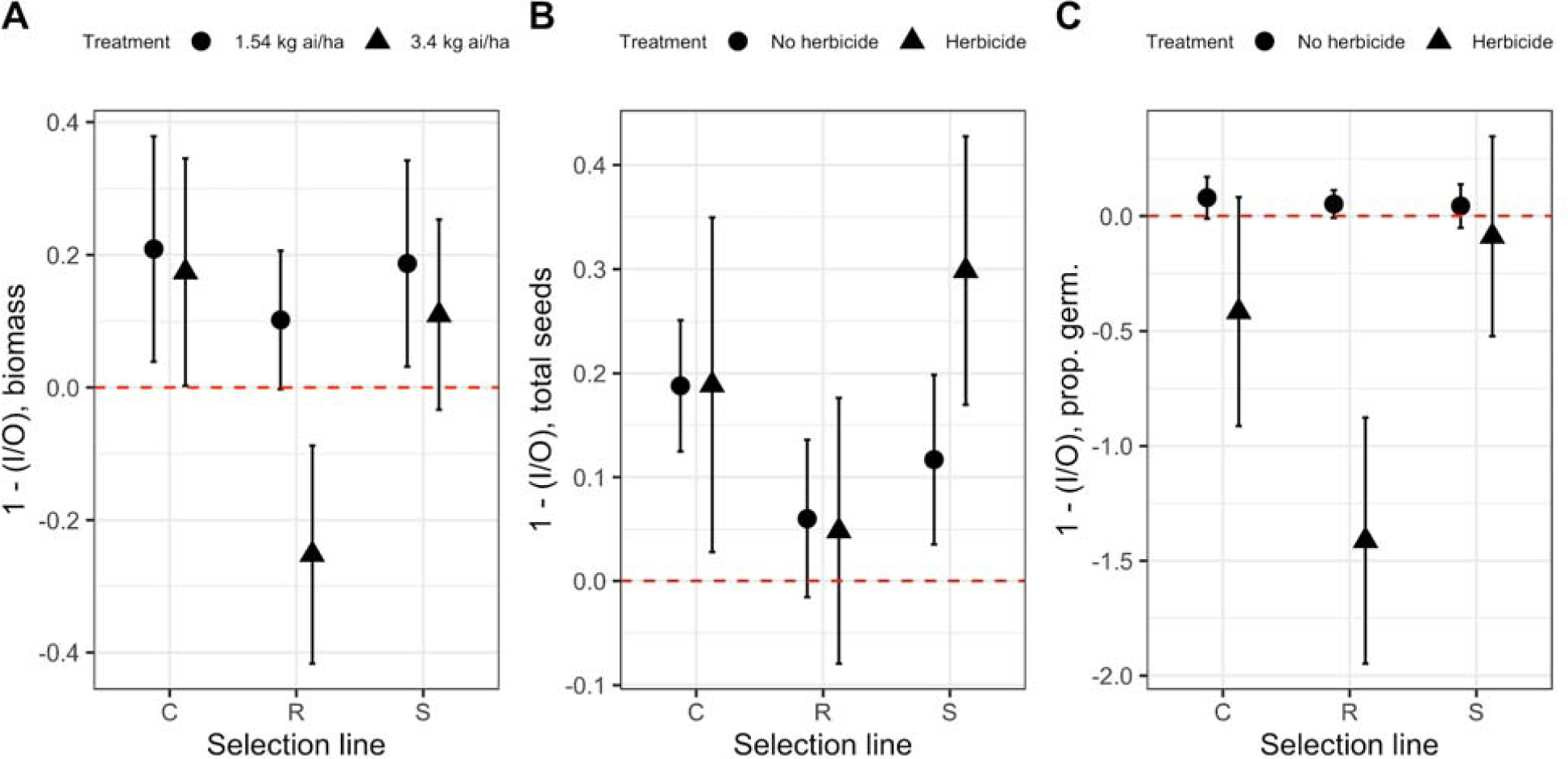
Inbreeding depression measured as ((Outcrossed - Inbred)/Outcrossed) for A) dried biomass, B) seed number, and C) proportion seed germination. Least square means for each family within selection lines and environments were used to determine averages across selection lines/environments for biomass and total seeds while means were used for germination (± standard error).

### Gene expression differs according to selection line, breeding history, and their interaction in the presence and absence of herbicide

We created groups of genes with correlated expression patterns (called modules) using weighted correlation network analysis, and then determined which of these modules had expression differences that correlated with the levels of each factor of interest (*i*.*e*. susceptible vs control vs resistant lines; outcrossed vs inbred). The number of modules that were created differed depending on the analysis, ranging from 23-88 (Table S6). The number of genes per module ranged from 21 to 7942, with the median ranging from 58 to 133 (Table S6).

Selection line was correlated with six to 13 modules depending on the environment and collection time. In the non-herbicide environment, selection line was correlated with 13 different modules that together contain 1107 genes, showing that the artificially evolved resistant and susceptible lines exhibited different patterns of gene expression even in the absence of herbicide (p-value range = <0.0001-0.03). More specifically, compared to the susceptible lines, resistant lines exhibited higher expression for modules that were enriched in GOterms including glycolytic processes, protein phosphorylation, defense response and amylopectin biosynthetic process, and exhibited lower expression for modules with GO terms including protein autophosphorylation, defense responses, regulation of cellular pH and protein phosphorylation (Table S7).

In the presence of herbicide, seven modules (containing 10,740 genes) were correlated with selection line at eight hours post-herbicide application (p-value range = 0.002-0.04) whereas six modules (containing 406 genes) were correlated with selection line at 32 hours post-herbicide application (p-value range < 0.0001-0.05; Table S6). At eight hours post-herbicide application the resistant lines, compared to susceptible lines, had higher expression for modules that were enriched in GOterms including regulation of cellular protein metabolic processes, photoperiodism and regulation of DNA replication, and lower expression for modules with GO terms including transport, defense response and cellular protein modification process (Table S7). At 32 hours post-herbicide application the resistant lines, again compared to susceptible lines, had higher expression for modules enriched in GOterms including regulation of protein complex disassembly and lower expression for modules with GO terms including cellular protein modification process, photoperiodism and cellular response to red or far red light (Table S7).

We next examined the potential for differences in gene expression between individuals produced by inbreeding or outcrossing (breeding type effect). We first identified modules that correlated with breeding type separately within the non-herbicide environment and within the herbicide present environment at two different time points (eight hours and 32 hours post-herbicide). We found that only the non-sprayed plants exhibited any module that correlated with breeding type (p-value = 0.03): the inbred plants exhibited higher expression for the genes correlated in one module (contained 68 genes), which was enriched for GOterms related to intracellular protein transport/localization (Table S7).

We further examined the potential for differences in gene expression according to breeding type (*i*.*e*. inbred versus outcrossed individuals) within selection lines. We again did so separately within the non-herbicide environment and in the presence of herbicide at two different time points (eight hours and 32 hours post-herbicide). In the presence of herbicide, we found different correlation patterns between breeding type and gene expression modules within the control and susceptible selection lines compared to that of the resistant selection lines (Table S6; Fig 5A and B). Specifically, at eight hours post herbicide application, both the susceptible and the control selection lines had modules that were significantly correlated with breeding type (p-value range = 0.01-0.03). In these modules, genes were up-regulated in inbred compared to outcrossed plants (1 and 4 modules, respectively; Fig 5A) suggesting that early responses of inbred and outcrossed plants differed in control and susceptible lines. These modules contained genes involved in organophosphate catabolic processes and chloroplast location (susceptible selection line, not significantly enriched), carbohydrate metabolic process, cell wall organization or biogenesis and cellular protein modification process (significantly enriched in the control line; Table 1; Table S7).

**Table 1.** The most significant GO terms for each module identified in the WGCNA that was significantly correlated with breeding type.

| Group | Module | P-value | Go Term |
| --- | --- | --- | --- |
| 8-hour post spray | saddlebrown | 1.30E-01 | Nucleotide catabolic process |
|  |  | 1.30E-01 | Chloroplast relocation |
|  |  | 1.30E-01 | Chloroplast localization |
|  |  | 1.30E-01 | Organophosphate catabolic process |
|  |  | 1.30E-01 | Plastid localization |
|  |  | 1.30E-01 | Establishment of plastid localization |
|  |  | 1.30E-01 | Nucleoside phosphate catabolic process |
|  | brown | 9.10E-08 | Carbohydrate metabolic process |
|  |  | 9.10E-08 | Cell wall organization or biogenesis |
|  | purple | 4.90E-13 | Cell wall organization or biogenesis |
|  | lightgreen | 9.10E-08 | Carbohydrate metabolic process |
|  |  | 9.10E-08 | Cell wall organization or biogenesis |
|  | lightsteelblue1 | 1.30E-02 | Cellular protein modification process |
|  |  | 1.30E-02 | Protein phosphorylation |
|  |  | 1.30E-02 | Phosphorus metabolic process |
|  |  | 1.30E-02 | Phosphate-containing compound metabolic process |
|  |  | 1.30E-02 | Protein modification process |
|  |  | 1.30E-02 | Protein autophosphorylation |
|  |  | 1.30E-02 | Regulation of protein serine/threonine kinase activity |
| 32-hour post<br>spray | saddlebrown | 7.70E-03 | Cation transport |
|  |  | 7.70E-03 | Nicotianamine metabolic process |
|  |  | 7.70E-03 | Nicotianamine biosynthetic process |
|  |  | 7.70E-03 | Tricarboxylic acid biosynthetic process |
|  | tan | 1.70E-02 | Response to UV-B |
|  | sienna3 | 2.10E-14 | Translation |
|  |  | 2.10E-14 | Peptide biosynthetic process |

**Fig 5.**
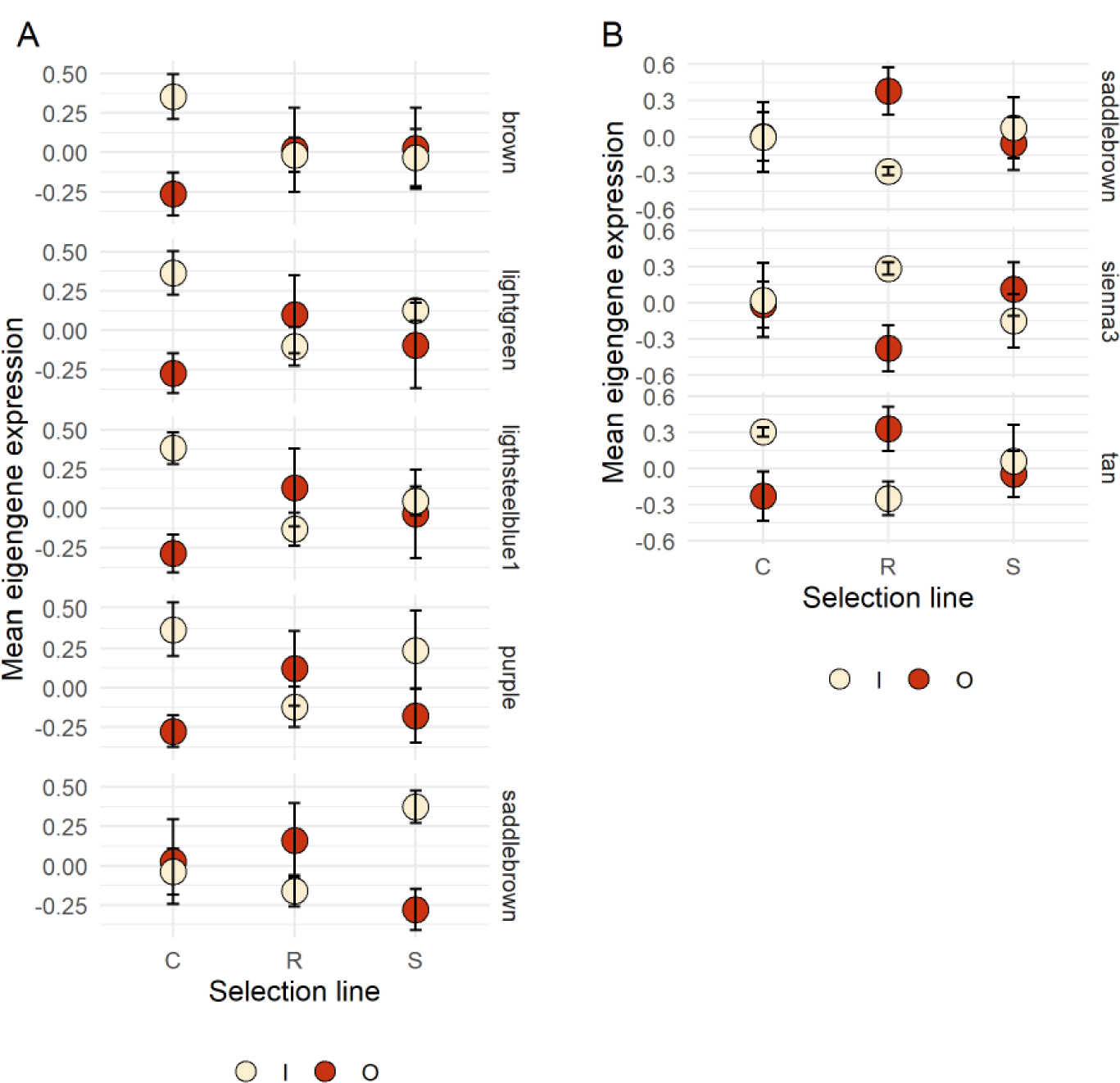
Mean eigengene expression (± standard error) by breeding type and selection line for sprayed 8 hour (A) and sprayed 32 hour (B). Only WGCNA modules (named by color) that were significantly different (p-value range = 0.01-0.05) by breeding type in at least one of the selection lines are shown.

Comparatively, for the samples collected 32-hours post herbicide, only the resistant selection line exhibited gene expression modules that were significantly correlated with breeding type (p-value range = 0.01-0.05). Of the three modules that were correlated with breeding type, two of these showed downregulation in inbred compared to outcrossed individuals (Fig 5B). Genes within these two modules were involved in cation transport and responses to UV-B (Table 1; Table S7). Interestingly, one module was upregulated in selfed compared to outcrossed resistant individuals, and it included genes involved in translation and peptide biosynthetic processes. These results suggest that at later time points, inbred resistant plants continued performing DNA replication and perhaps growth compared to resistant outcrossed plants. In the absence of herbicide, no modules correlated with breeding type within selection lines.

## Discussion

In this work, we examined the potential for inbreeding depression in lineages selected for divergent levels of herbicide resistance. Previous work in this species identified a correlation between the level of resistance and the mating system, such that more resistant populations exhibited a higher rate of selfing (Kuester et al., 2017). However, this association might not be maintained over time if resistant individuals exhibit inbreeding depression, or a cost associated with selfing. Thus, our aim was to determine if inbreeding depression was apparent in lines selected to exhibit higher levels of herbicide resistance in order to make longer-term predictions about the previously identified association between resistance and the mating system. We used a combination of growth chamber and field experiments to examine a variety of resistance and fitness related traits, and we paired this work with an examination of gene expression to identify effects associated with breeding history, herbicide resistance, and their interaction on the transcriptome.

Similar to Debban et al., (2015), our growth chamber experiment showed that the selection lines varied in the level of resistance; individuals selected for increased resistance survived herbicide application at a higher proportion and showed greater biomass post-application compared to lineages selected for decreased resistance (at twice the recommended field dose). We found evidence of inbreeding depression in biomass in the growth chamber and flower number and seed number in the field. Interestingly, inbreeding depression was often non-existent for individuals from the increased resistance lines, especially when measured in the presence of herbicide. Strikingly, in the increased resistance lines, we identified outbreeding depression in both biomass and seed germination in the presence of herbicide. Further, at the transcriptome level, the resistant selection lines had differing patterns of gene expression according to breeding type (inbred vs outcrossed) compared to the control and susceptible selection lines. These data show that the expression of inbreeding depression varied among selection lines -- both across fitness related traits and at the level of the transcriptome. That we uncovered outbreeding depression in the presence of herbicide in the resistant lines across two separate experiments and across different fitness-related traits suggests that resistance and a selfing mating system may remain associated in populations experiencing herbicide application. Below, we discuss the implications of this result for applied resistance evolution and more broadly for mating system evolution.

### The expression of inbreeding depression varies among artificially evolved resistance lines

Many of the traits we examined in this work exhibited inbreeding depression. In the controlled conditions of the growth chamber, we found inbreeding depression in both biomass and survival post-herbicide; in the field, we found evidence for inbreeding depression across most fitness related traits in both the control and herbicide-present environments. Thus, our work shows that inbreeding depression is evident at different stages of the *I. purpurea* life cycle. Interestingly, however, we found no evidence to suggest that the expression of inbreeding depression is greater in more stressful environments (in our case, the presence of herbicide), a finding that differs from many other examples (Fox & Reed, 2011). We identified one notable exception across traits: we uncovered significant inbreeding depression for the total number of seeds produced by the susceptible selection lines in the presence but not absence of herbicide (Fig 4B). In other plants, such as *Sabatia angularis (Dudash, 1990)* and *Lobelia* (Johnston, 1992), inbreeding depression is more pronounced in the more stressful conditions of the field compared to the greenhouse, and inbreeding depression is likewise more pronounced in plants that experience competition (Cheptou et al., 2000; Schmitt & Ehrhardt, 1990), drought (Hauser & Loeschcke, 1996), nutrient stress (Helenurm & Schaal, 1996), and damage from herbivores (Carr & Eubanks, 2002; Du et al., 2008; Hull-Sanders & Eubanks, 2005; Ivey & Carr, 2012; Stephenson et al., 2004). Thus, while not a primary focus of the work presented here, our results do not consistently or strongly support the idea that increased stress leads to higher values of inbreeding depression.

The main focus of our work -- assessing whether or not the resistant selection lines exhibited inbreeding depression -- led to unexpected findings. Across both the growth chamber and field experiment, we found little evidence of inbreeding depression in the increased resistance selection lines, and even found evidence of outbreeding depression in two traits -- biomass at the highest dose of herbicide in the growth chamber and in seed germination in the presence of herbicide in the field. In fact, resistant, inbred individuals showed 1.8x higher germination of their seeds than resistant, outcrossed individuals, suggesting that under sprayed conditions, selection would favor increased selfing. If this result from a single experimental population is applicable to the dynamic across natural populations, it suggests that the previously identified relationship between high resistance and a highly selfing mating system in this species (Kuester et al., 2017) may be maintained over time.

Strikingly, these results -- a lack of inbreeding depression or evidence of outbreeding depression in herbicide adapted lineages -- are similar to results from a study of inbreeding depression in salt and cadmium adapted Drosophila lineages. To investigate the expression of inbreeding depression in different environments, (Long et al., 2013) reared replicate populations of *D. melanogaster* on diets including either CdCl_2_ or NaCl for >20 generations. Populations adapted to CdCl_2_ or NaCl exhibited much lower inbreeding depression when exposed to that particular stressor but higher inbreeding depression when exposed to a novel or mismatched environments -- *i*.*e*., when CdCl_2_ or NaCl adapted flies were exposed to NaCl or CdCl_2_, respectively. The explanation for this result is that past selection likely reduced the level of polymorphism at the sites under selection, leading to a negative covariance between the selection coefficient and amount of polymorphism and hence a lower value of inbreeding depression when populations were exposed to their adapted environment (Yun & Agrawal, 2014). Such an effect seems a likely explanation for our result of lower and often nonsignificant values of inbreeding depression in our increased resistance lines and potentially even serves as an explanation for the presence of outbreeding depression in adapted lines when exposed to herbicide.

Additionally, underdominance or epistatic interactions among loci are potential explanations for this result. If loci tightly linked to sites under selection exhibited underdominance (*i*.*e*. a disadvantage in heterozygous compared to homozygous form), this could lead to lower fitness of outcrossed resistant individuals compared to inbred resistant individuals and hence the pattern of outbreeding depression we uncovered in some traits. Further, if the loci underlying adaptation to herbicide exhibit negative interactions with alleles of loci from genetic backgrounds introduced *via* outcrossing, such Dobzhansky-Muller like epistatic interactions (Bikard et al., 2009; Bomblies et al., 2007) could lead to lower fitness of resistant, outcrossed individuals compared to resistant, inbred individuals. Finally, based on previous work from this and other natural populations, it is possible that higher resistance in these artificially evolved lineages is polygenic (Leslie & Baucom, 2014; Van Etten et al., 2020). Because inbreeding reduces recombination, individuals that were outcrossed may have inherited combinations of alleles that provide less resistance compared to inbred types. Further research will be necessary to delineate between these possibilities.

### Gene expression changes parallel organismal-level traits

To examine the effect of resistance, breeding history, and the interaction between breeding history and resistance on gene expression, we examined groups of similarly expressed genes using a network based analysis. Using this approach, we found that the expression of many genes differed among the selection lines, especially soon after spraying. Interestingly, breeding history led to few gene expression differences. However, and similar to our phenotypic results, inbreeding had a different effect on gene expression in the resistant selection line compared to the control and susceptible lines.

The selection lines differed in gene expression across many genes, especially at the earliest time point examined (eight hour post-spray). At eight hours after spraying, 10,740 genes (37%) were in a module with differential expression between the selection lines, suggesting that the resistant individuals responded early and differently than the other lines following application of the herbicide. At this time point, the resistant lines upregulated growth-related processes and downregulated defense-related genes. By 32 hours post-herbicide, however, only 406 genes were in modules that showed differential regulation between selection lines, perhaps suggesting that the majority of difference in resistance may happen early (a result similar to that found in (Kohlhase et al., 2019). We previously found genomic regions showing signatures of selection in multiple resistant populations (Van Etten et al., 2020) that were enriched for genes involved in detoxification - cytochrome P450s, ABC transporters, glycosyltransferases, and glutathione S-transferases. Two of the modules identified here (“blue” and “turquoise”) had many of these genes, with the resistant lines showing lower expression than the susceptible lines (Table S8). Studies in two other glyphosate resistant species (*Conyza bonariensis* and *Lolium multiflorum*) have also found differential expression in detoxification genes, but with a tendency for these genes to have higher expression in resistant lines (Cechin et al., 2020; Piasecki et al., 2019).

Surprisingly, examining gene expression between inbred and outcrossed individuals over all the treatments showed very few differences. This could be due to a variety of reasons. Inbreeding depression is generally thought to be due to the action of many slightly deleterious genes when in homozygous form (Charlesworth & Charlesworth, 1999; Latter, 1998) and therefore many more individuals than assayed here would be needed to identify differences. Further, the genes involved in inbreeding depression may vary between individuals, such that different individuals are likely to have different sets of genes in the recessive homozygous state, meaning that we would not necessarily expect consistent changes associated with inbreeding depression across all individuals (Carr & Dudash, 2003). Finally, the expression of inbreeding depression may be more extreme in other tissues, such as tissues in the embryo or in seed production, rather than the leaves that we sampled. While there are no studies, to our knowledge, that compare inbreeding depression at the transcriptomic level across different plant tissues, different tissues do show different expression profiles (Schmid et al., 2005), and thus may show a differential effect associated with inbreeding.

Perhaps for some of the above reasons, little work on the transcriptomic effects of inbreeding has been performed. Much of the work has been done in *Drosophila melanogaster* in which inbreeding consistently leads to upregulation of genes involved in metabolism, stress and defense responses (Ayroles et al., 2009; García et al., 2012, 2013; Garcia et al., 2013; Kristensen et al., 2005, 2006). Other studies have shown that inbreeding reduces expression of stress response genes (Franke & Fischer, 2015; Kariyat et al., 2012), whereas others do not find a clear pattern (Zhao et al., 2019) or show no difference in expression (Hansson et al., 2014). In our work, the differences we find associated with inbreeding depression are in genes involved in protein transport and localization, which is different from these examples. Clearly, the causes and consequences of inbreeding on gene expression are in need of further research.

Interestingly, despite few overall differences due to breeding history, we found differences according to breeding type among the selection lines. At the 8-hour post spray, both the susceptible and control lines showed an effect of breeding type while the resistant lines exhibited similar expression between outcrossed and inbred individuals. Comparatively, at 32 hours post spray, selfed and outcrossed individuals exhibited different expression patterns only in the resistant lines. At this time point, the resistant inbred plants downregulate nicotianamine metabolic process related genes, which play a role in the uptake of metal from the soil, along with metal ion transport related genes. Since glyphosate is known to chelate several metals (Cakmak et al., 2009), these genes may play a role in the differing response of inbred and outcross plants. The inbred individuals also show increased expression of several genes involved in abiotic stress tolerance (in the “sienna3” module; Table S7), which may play a role in mitigating the stress induced by the herbicide treatment. The inbred individuals are also upregulating genes involved in translation. Glyphosate inhibits the shikimate pathway, which leads to a shortage of aromatic amino acids, which should cascade into suppressing translation (Faus et al., 2015; Hinnebusch, 2005; Lageix et al., 2008). The upregulation of translation-related genes may show that inbred plants are better able to avert the shutdown of translation. Interestingly, the inbred individuals exhibited lower expression of photoperiod-related genes, which is the same set of genes we found were down regulated in the resistant lines compared to the susceptible lines. This suggests that the inbred resistant individuals have a different response than the outcrossed resistant individuals, a finding that parallels the organismal-level patterns we see in the fitness-related traits.

## Conclusion

Across the experiments and traits measured, we found that the resistant selection line shows little to no inbreeding depression among the examined traits and even an advantage to inbreeding under sprayed conditions. Thus, over time and given continued herbicide application, we would expect that the previously identified relationship between high resistance and selfing across natural populations may be maintained in this species. This association could have repercussions for the evolution and maintenance of resistance within populations and thus influence how we manage resistance in this and other weeds that employ mixed-mating systems. For example, if adaptation to herbicide leads to populations with a high rate of selfing, relying on gene flow from susceptible populations as an important source of susceptible alleles would not be a viable strategy for the mitigation of resistance (Maxwell et al., 1990). This potential control tactic is rarely used, however. In practice, when a population evolves herbicide resistance, a different herbicide is suggested for control, and in that light, it is unknown how inbred lines resistant to one herbicide will respond relative to outcrossed resistant lines when exposed to a novel herbicide. If the results from NaCl and CdCl_2_ adapted *Drosophila (Long et al*., *2013)* are an appropriate proxy, and assuming some level of cross-resistance between herbicides, it would suggest that inbred, resistant lineages should do worse than outbred, resistant lineages when exposed to a new herbicide -- a result that would align with theoretical predictions that strong selection should lead to an outcrossing, resistant population. Future experimentation exposing herbicide adapted, inbred individuals to different herbicide modes of action is thus warranted, as is the generality of an association between herbicide resistance and the mating system across more plant species, like that identified in (Kuester et al., 2017).

Reproductive assurance from increased self-fertility can heighten the likelihood that a population will survive environmental change so long as the gain in seed productivity from selfing is not outweighed by inbreeding depression (Levin, 2012). The findings reported here raise important questions as to whether plant species adapting to other strong agents of selection may likewise exhibit changes in the level of inbreeding depression, thus influencing (or reinforcing) mating system evolution. Beyond the dynamic that we examine -- i.e., agricultural weed populations subjected to population bottlenecks from herbicide application -- our results are likely applicable to a variety of scenarios in which populations of self-compatible species are rapidly reduced in size. While mating system changes have been found in species adapted to heavy-metal and serpentine soils (e.g. *Thlaspi caerulescens, Anthoxanthum odoratum, Lasthenia* and *Mimulus* sp; (Antonovics et al., 1971; Dubois et al., 2003; Macnair & Gardner, 1998), the potential for inbreeding depression has not, to our knowledge, been examined in these systems. Our study is also likely applicable to plant populations experiencing population size reductions due to climate change, whether as a consequence of changing climatic conditions (drought, early frosts) or from existing in small populations on the trailing edge of a species’ range. Future research examining the potential for mating system evolution and inbreeding depression in lineages adapting to strong selective agents will be necessary to determine the ubiquity of the findings reported here.

## Acknowledgements

We thank A. Jankowiak for performing crosses and generating seeds, D. York, and T. Marrs for maintaining the field experiment, collecting data and counting seeds, M. Palmer and R. Grese for providing support at the Matthaei Botanical Gardens, K. Johnson, S. Paranjape and T. Newsum for assistance with the growth chamber experiment, and E. Josephs for providing comments on an earlier manuscript draft. This work was funded by the University of Michigan and USDA NIFA (United States Dept of Agriculture, NIFA: https://nifa.usda.gov/) (awards 24892 & 28497).

## Author Contributions

RSB and MVE designed the research, MVE and AS performed the growth chamber and field studies, MVE and RSB analyzed the gene expression and phenotype data, and RSB and MVE wrote the paper.

## Data accessibility

Data will be made available upon acceptance (raw sequencing data and transcriptome will be archived in NCBI)

## Supplementary Information

**STable 1.** Analysis of deviance table from a logistic regression examining the proportion of individuals alive versus dead/almost dead across both rates of herbicide (Overall) and within each treatment in the growth chamber study.

| Overall | Deviance | df | P-value |
| --- | --- | --- | --- |
| Treatment | 62.54 | 1 | **<0.001** |
| Breeding type | 4.96 | 2 | **0.03** |
| Selection line | 29.81 | 2 | **<0.001** |
| Within treatment |  |  |  |
| 1X herbicide (1.54 kg ai/ha) | Deviance | df | P-value |
| Selection line | 13.39 | 2 | **0.001** |
| Breeding type | 2.33 | 1 | 0.126 |
| 2X herbicide (3.4 kg ai/ha) | Deviance | df | P-value |
| Selection line | 16.864 | 2 | **<0.001** |
| Breeding type | 2.691 | 1 | 0.1 |

**STable 2.** Mixed model analysis of variance examining the effect of treatment, breeding type, selection line, their interactions, and maternal line on the dried biomass of individuals remaining post-herbicide application in the growth chamber study. Individual dried biomass was first standardized to average maternal line dried biomass from the control prior to analysis.

|  | <u>Standardized biomass</u> |  |  |  |
| --- | --- | --- | --- | --- |
| *Fixed effects* | Num df | Den df | F-value | P-value |
| Treatment | 1 | 495.47 | 14.82 | **<0.001** |
| Breeding type | 1 | 498.59 | 6.87 | **0.009** |
| Selection line | 2 | 14.15 | 2.96 | 0.084 |
| Breeding type*Selection line | 2 | 498.28 | 3.23 | **0.040** |
| Breeding type*Treatment | 1 | 495.47 | 3.96 | **0.047** |
| *Random effects* | | df | $\eta^2$ | P-value |
| Maternal line |  | 1 | 29.84 | **<0.001** |
| <u>1X Herbicide</u> | Num df | Den df | F value | P-value |
| Selection line | 2 | 15.34 | 1.66 | 0.222 |
| Breeding type | 1 | 236.47 | 7.17 | **0.008** |
| Breeding type*Selection line | 2 | 236.26 | 0.235 | 0.791 |
| | df | | $\eta^2$ | P-value |
| Maternal line | 1 |  | 8.53 | **0.003** |
| <u>2X Herbicide</u> | Num df | Den df | F value | P-value |
| Selection line | 2 | 14.1 | 4.31 | **0.035** |
| Breeding type | 1 | 245.67 | 0.42 | 0.517 |
| Breeding type*Selection line | 2 | 245.38 | 4.59 | **0.011** |
|  | df |  | Chisq | P-value |
| Maternal line | 1 |  | 9.75 | **0.002** |

**STable 3.** Mixed model analysis of variance examining the effect of breeding type (inbred/outcrossed), selection line (R/S/C), treatment (herbicide, control), block, and their interactions, along with the effect of maternal line, on flowering day, flower number, seed number, and germination of seeds produced by individuals in the field experiment.

|  | <u>Flowering day</u> |  |  | <u>Flower number</u> |  |  | <u>Seed number</u> |  |  | <u>Seed germination</u> |  |  |
| --- | --- | --- | --- | --- | --- | --- | --- | --- | --- | --- | --- | --- |
| *Fixed effects* | Num, den df | F-value | p-value | Num, den df | F-value | p-value | Num, den df | F-value | p-value | df | $\eta^2$ | p-value |
| Breeding type | 1, 1405.05 | 2.82 | 0.09 | 1, 1412 | 11.27 | **<0.001** | 1, 1365.47 | 8.56 | **0.003** | 1 | 1.51 | 0.22 |
| Selection line | 2, 19.58 | 0.61 | 0.55 | 2, 26.04 | 0.19 | 0.83 | 2, 19.64 | 0.09 | 0.91 | 2 | 0.26 | 0.88 |
| Treatment | 1, 1405.09 | 8,085.08 | **<0.001** | 1, 1412 | 8.81 | **<0.01** | 1, 1365.47 | 1248.48 | **<0.001** | 1 | 390.39 | **<0.001** |
| Block | 1, 1405.03 | 0.39 | 0.53 | 1, 1412 | 2.22 | 0.14 | 1, 1365.47 | 0.03 | 0.84 | 1 | 0.35 | 0.56 |
| Breeding type*Selection line | 2, 1405.05 | 0.09 | 0.92 | 2, 1412 | 0.84 | 0.42 | 2, 1365.47 | 1.83 | 0.16 | 2 | 1.47 | 0.48 |
| Breeding type*Treatment | 1, 1405.05 | 1.38 | 0.24 | 1, 1412 | 3.82 | **0.05** | 1, 1365.47 | 2.05 | 0.15 | 1 | 7.74 | **<0.01** |
| Selection line*Treatment | 2, 1405.09 | 1.31 | 0.27 | 2, 1412 | 0.91 | 0.4 | 2, 1365.47 | 0.03 | 0.97 | 2 | 1.08 | 0.58 |
| Breeding Type*Selection line*Treatment | 2, 1405.05 | 0.23 | 0.79 | 2, 1412 | 0.82 | 0.44 | 2, 1365.47 | 1.77 | 0.17 | 2 | 5.35 | 0.07 |
| *Random effects* | df | $\eta^2$ | p-value | df | $\eta^2$ | p-value | df | $\eta^2$ | p-value | | | |
| Maternal line | 1 | 0.016 | 0.9 | 1 | 0.016 | 0.9 | 1 | 7.44 | **<0.01** | -- | -- | -- |

**STable 4.** Mixed model analysis of variance examining the effect of breeding type (inbred/outcrossed), selection line (R/S/C), block, and their interactions, along with the effect of maternal line, on flowering day, flower number, seed number, and germination of seeds produced by individuals in either the control (no herbicide) or herbicide environment in the field experiment.

| <u>Control</u> | <u>Flowering day</u> |  |  |  | <u>Flower number</u> |  |  |  | <u>Seed number</u> |  |  |  |
| --- | --- | --- | --- | --- | --- | --- | --- | --- | --- | --- | --- | --- |
| *Fixed effects* | Num df | Den df | F-value | P-value | Num df | Den df | F-value | P-value | Num df | Den df | F-value | P-value |
| Breeding type | 1 | 696.01 | 8.64 | **0.003** | 1 | 698.00 | 1.25 | 0.265 | 1 | 698.00 | 7.33 | **0.007** |
| Selection line | 2 | 19.50 | 1.13 | 0.343 | 2 | 23.47 | 0.13 | 0.877 | 2 | 24.63 | 0.13 | 0.877 |
| Block | 1 | 669.22 | 3.98 | **0.046** | 1 | 698.00 | 4.15 | **0.042** | 1 | 698.00 | 5.28 | **0.022** |
| Breeding type*Selection line | 2 | 696.01 | 0.58 | 0.551 | 2 | 698.00 | 0.14 | 0.867 | 2 | 698.00 | 0.54 | 0.584 |
| *Random effects* | | df | $\eta^2$ | P-value | | df | $\eta^2$ | P-value | | df | $\eta^2$ | P-value |
| Maternal line |  | 1 | 7.18 | **0.007** |  | 1 | 0.09 | 0.759 |  | 1 | 0.21 | 0.645 |

| <u>Herbicide</u> | <u>Flowering day</u> |  |  |  | <u>Flower number</u> |  |  |  | <u>Seed number</u> |  |  |  |
| --- | --- | --- | --- | --- | --- | --- | --- | --- | --- | --- | --- | --- |
| *Fixed effects* | Num df | Den df | F-value | P-value | Num df | Den df | F-value | P-value | Num df | Den df | F-value | P-value |
| Breeding type | 1 | 693.06 | 0.09 | 0.761 | 1 | 698 | 12.03 | **<0.001** | 1 | 651.81 | 6.14 | **0.013** |
| Selection line | 2 | 19.10 | 0.44 | 0.649 | 2 | 22.34 | 0.60 | 0.556 | 2 | 19.57 | 0.04 | 0.958 |
| Block | 1 | 693.04 | 0.16 | 0.689 | 1 | 698 | 13.07 | **<0.001** | 1 | 651.51 | 1.96 | 0.162 |
| Breeding type*Selection line | 2 | 693.06 | 0.03 | 0.976 | 2 | 698 | 1.32 | 0.269 | 2 | 651.80 | 2.25 | 0.106 |
| *Random effects* |  | df | Chisq | P-value |  | df | Chisq | P-value |  | df | Chisq | P-value |
| Maternal line |  | 1 | 9.06 | **0.003** |  | 1 | 1.42 | 0.233 |  | 1 | 7.66 | **0.006** |

|  | <u>Proportion seeds germinated</u> |  |  |  |  |  |
| --- | --- | --- | --- | --- | --- | --- |
|  | No herbicide |  |  | Herbicide |  |  |
| | df | $\chi^2$ | P-value | df | $\chi^2$ | P-value |
| Breeding type | 1 | 5.451 | **0.020** | 1 | 3.889 | **0.049** |
| Selection line | 2 | 0.714 | 0.700 | 2 | 0.787 | 0.675 |
| Block | 1 | 0.000 | 0.992 | 1 | 1.545 | 0.214 |
| Breeding type*Selection line | 2 | 0.286 | 0.867 | 2 | 6.733 | **0.035** |

**STable 5.** Least square means (standard errors) of fitness related traits examined in the field experiment by treatment, selection line, and breeding type.

| Treatment | Selection Line | Breeding type | Flowering day | Number flowers | Number seed | Prop. seed germinated |
| --- | --- | --- | --- | --- | --- | --- |
| Herbicide absent | Control | Inbred | 60.08 (0.44) | 7.81 (0.66) | 2180 (200.02) | 0.57 (0.02) |
|  |  | Outbred | 59.69 (0.31) | 8.58 (0.41) | 2695 (120.36) | 0.63 (0.01) |
|  | Resistant | Inbred | 60.38 (0.44) | 8.27 (0.66) | 2280 (200.02) | 0.60 (0.01) |
|  |  | Outbred | 59.60 (0.31) | 8.51 (0.41) | 2464 (120.36) | 0.64 (0.01) |
|  | Susceptible | Inbred | 61.07 (0.44) | 8.29 (0.66) | 2275 (200.02) | 0.60 (0.02) |
|  |  | Outbred | 59.94 (0.31) | 8.68 (0.41) | 2635 (120.36) | 0.64 (0.01) |
| Herbicide present | Control | Inbred | 88.95 (1.14) | 9.03 (0.82) | 310 (48.88) | 0.37 (0.05) |
|  |  | Outbred | 88.39 (0.83) | 10.54 (0.52) | 366 (31.73) | 0.30 (0.01) |
|  | Resistant | Inbred | 87.69 (1.14) | 8.84 (0.82) | 274 (49.27) | 0.49 (0.05) |
|  |  | Outbred | 87.53 (0.83) | 9.76 (0.52) | 303 (31.72) | 0.27 (0.01) |
|  | Susceptible | Inbred | 88.32 (1.14) | 7.52 (0.82) | 267 (49.69) | 0.27 (0.04) |
|  |  | Outbred | 88.09 (0.83) | 10.43 (0.52) | 396 (31.66) | 0.30 (0.01) |
| IBD, no herbicide |  |  | 0.013 | 0.055 | 0.14 | 0.075 |
| IBD, herbicide |  |  | -0.003 | 0.174 | 0.201 | -0.308 |
| IBD, overall |  |  | 0.007 | 0.114 | 0.17 | -0.116 |

STable 6. The significantly enriched GOTerms for each significant module. The Effect column shows the comparion in which the module was significant. The Module column is the name of the module. The correlation coefficient is the in Correlation column and the associated p-value is in the Correlation p-value column. The GoTerm enrichment FDR value is in the Enrichment FDR column followed by the number of genes in the module that had that GOTerm and the total number of genes in Arabidopsis that have that GOTerm. Lastly the GOTerm (biological processes only) are in the last column. Link to table <u>here</u>.

**STable 7.** Information about the modules and genes in each comparison. The comparison is in the Effect column. The 2nd column shows the number of significant modules and the total number of modules. The third column shows the total number of genes in the significant modules and the total number of genes that were assigned to a module (many genes were unassigned in some analyses). The final column calculates the proportion of the total assigned genes that fell into a significant module.

| Effect | # sig mods/ttl mods | #genes / ttl genes in mods | prop of genes that differ |
| --- | --- | --- | --- |
| Selection line, NS (S, C, R) | 13/51 | 1107/18003 | 0.06148975171 |
| Selection line, S8 (S, C, R) | 7/61 | 10740/29243 | 0.3672673802 |
| Selection line, S32 (S, C, R) | 6/88 | 406/18855 | 0.02153274993 |
| Breeding type, NS (O, I) | 1/51 | 68/18003 | 0.003777148253 |
| Breeding type, S8 (O, I) | 0/61 | 0/29243 | 0 |
| Breeding type, S32 (O, I) | 0/88 | 0/18855 | 0 |
| Breeding type within line, NS, (O, I; S, C, R) | 0,0,0/23 | 0,0,0/5754 | 0 |
| Breeding type within line, S8, (O, I; S, C, R) | 1,4,0/49 | 93,1153,0/9097 | 93,1153,0/9097 |
| Breeding type within line, S32, (O, I; S, C, R) | 0,0,3/86 | 0,0,416/11588 | 0,0,416/11588 |

STable 8. Gene annotations for genes contained in signficant modules for each of the effects tested. The first column shows the comparison where the module was signficant. The second column is the transcript ID. The third column is the module name. The fourth coulmn is the group in which there was a difference between levels of the effect. The fifth column is the Arabidopsis gene ID, followed by the gene name and description. The last set of columns are the GOTerms and their descriptions. Link to table <u>here</u>.

